# DeepRank-Ab: a scoring function for antibody-antigen complexes based on geometric deep learning

**DOI:** 10.64898/2025.12.03.691974

**Authors:** Xiaotong Xu, Ilaria Coratella, Victor Reys, Alexandre MJJ Bonvin

## Abstract

Gaining structural insights into the interactions between antibodies and their corresponding antigens is essential for understanding immune recognition and for guiding therapeutic antibody design. However, accurately modelling these complexes remains a significant challenge for both physics-based docking approaches and AI-based, co-folding methods such as AlphaFold3. These methods not only struggle to generate near native conformations, but, more critically, they often fail to score and rank such conformations correctly, revealing fundamental limitations when applied to antibody-antigen systems.

To address these limitations, we present here DeepRank-Ab, a geometric deep learning-based scoring function tailored to the unique characteristics of antibody-antigen interfaces. Its development was enabled by a rigorously curated benchmark comprising more than 2.3 million decoys generated from 1,442 complexes, providing the diversity required for robust training and unbiased evaluation. Leveraging this resource, we systematically assessed multiple levels of graph representation, structural and energetic feature sets, and sampling strategies. Building on our previous DeepRank-GNN-esm work, our analysis identified that atom-level graph representation coupled with Voronoi-based surface decomposition and antibody-specific features is the most effective formulation for accurate scoring.

Across multiple independent test sets, including models from unbound unbound docking and structures generated by AlphaFold, DeepRank-Ab consistently outperforms all evaluated methods, including AF3, HADDOCK and state of the art scoring functions such as FTDMP. It increases the AF3 Top 1 success rate by 35.5% and improves the mean Top 1 DockQ by more than a factor of two. DeepRank-Ab further generalizes robustly beyond its training distribution, achieving 100% Top 5 success rate on external antibody-antigen CAPRI targets, surpassing all tested methods. Together, these results demonstrate that DeepRank-Ab is a highly effective scoring method that substantially improves the identification of near-native antibody-antigen conformations.

## 1. Introduction

Antibodies are protective proteins generated by B lymphocytes as part of the humoral immune response to antigens. Structurally, they are Y-shaped macromolecules composed of two heavy and two light chains, each containing constant and variable regions. The variable regions include the complementarity-determining regions (CDRs), which are hypervariable loops that form the antigen-binding site and determine antibody specificity and affinity. Sequence and structural variations within these loops give rise to the immense diversity of antibodies, estimated at 10¹⁵ to 10¹⁸ distinct specificities ^1,2^. Studying antibody-antigen interactions is therefore fundamental not only for understanding immune recognition but also for advancing vaccine and diagnostic development, as well as the design of therapeutic antibodies ^3^.

Modelling antibody-antigen interactions at atomic resolution remains highly challenging, even in the age of artificial intelligence. Co-folding algorithms ^4, 5^ rely heavily on co-evolutionary signals to infer residue-residue contacts, but such signals are largely absent in antibody-antigen pairs. Although AlphaFold3 (AF3) ^4^ reports success rates of 40-60% in modelling such complexes, these results are typically obtained only after extensive sampling and re-ranking using internal confidence metrics. Open-source reproductions of AF3, including OpenFold3 ^6^ and Boltz-2 ^7^ etc., still underperform compared to AF3 despite being trained on comparable data with similar architectures. Moreover, across all these approaches, a substantial gap exists between the best conformations produced by the models and those favoured in their rankings ^8^.

In AlphaFold2, model ranking relies on a weighted combination of ipTM and pTM ^5^ whereas AF3 additionally applies penalties for steric clashes and structural disorder ^4^. Several recent methods aim to improve ranking, including pDockQ2 ^9^ for chain-specific evaluation, actifpTM ^10^ for handling flexible regions, and ipSAE ^11^ for better separation of correct and incorrect interfaces. However, for antibody-antigen complexes, recent work ^12^ shows that the main source of error lies not in the ranking formula itself but in AlphaFold’s limited ability to estimate alignment error. This suggests that further refinement of AF3’s ranking scheme is insufficient, and that external scoring functions and/or improving AF3’s confidence prediction head ^13^ are needed to improve performance on these systems.

Given these challenges, physics-based modelling remains highly valuable. Accurate modelling of antibody-antigen interactions is largely dependent on obtaining a reliable starting structure—a task made difficult by the vast sequence diversity of antibodies and the intrinsic flexibility of their CDR loops ^14^. In particular, correct conformation of CDR-H3 have been shown to be essential for successful docking ^15^. The scoring problem is also severe in docking: only about 20% of the top-ranked models correspond to correct binding modes, reflecting the limited capability of current scoring functions, particularly in unbound docking scenarios where conformational changes occur during binding ^16^.

Deep learning approaches trained on docking decoys have been widely explored ^17–21^. Despite these advances, the scoring problem remains unsolved. Poor generalizability is a persistent issue. As shown by Stratiichuk *et al.*, scoring functions trained for rigid docking protocols often fail to generalize to decoys generated through flexible sampling^22^. One contributing factor is the limited diversity of available training data. Constructing large, heterogeneous docking benchmarks is difficult, and most scoring methods were trained on decoys produced by a single protocol and software. Data leakage poses a second challenge: strict separation between training and test sets is not always enforced, and common practices such as sequence-similarity or temporal filtering do not guarantee dataset independence ^23^, leading to inflated performance estimates.

To address this, we developed DeepRank-Ab, a deep-learning scoring function tailored specifically to antibody-antigen interfaces. Because of the lack of benchmark datasets covering both bound and unbound conformations while capturing the structural flexibility of antibody-antigen interactions, we first constructed a dataset of more than 2 million models generated from 1,442 antibody-antigen complexes under four docking protocols. Building upon our previous DeepRank-GNN-esm ^18^ framework, we systematically evaluated multiple levels of graphs, feature representations, and neural network architectures to optimize the overall model formulation. The final DeepRank-Ab models achieve performance that surpasses external methods, including AF3 ranking confidence ^4^, HADDOCK score ^24,25^, and one of the leading scoring function, FTDMP ^17^. DeepRank-Ab is available at https://github.com/haddocking/DeepRank-Ab.

## 2. Results

### 2.1 Assessment of AF3’s scoring capabilities

We first evaluated AF3’s ability to rank antibody-antigen complexes before developing DeepRank-Ab. From our benchmark test set, we selected 59 complexes released after the AF3 training cutoff date. For each complex, AF3 was run using 100 random seeds, generating a total of 500 models to ensure sufficient sampling.

AF3 achieves a Top 1 success rate of 40.7%, increasing to 59.3% as *K* increases, in line with reported values ^4^ (Figure 1a). However, both the Top 1 success rate and average Top K DockQ would be nearly doubled under the oracle setting which represents an optimal scoring scenario where the best model is ranked first. This observation highlights a central limitation of AF3’s scoring: the method fails to recognize acceptable models and fail to prioritize high-quality structures. Figure 1b illustrates this issue with an example (PDBID: 7PS0). AF3 generates a high-quality model (DockQ = 0.6) but ranks it at 498^th^, while the Top-ranked model fails to form the correct contacts with antigen.

**Figure 1.**
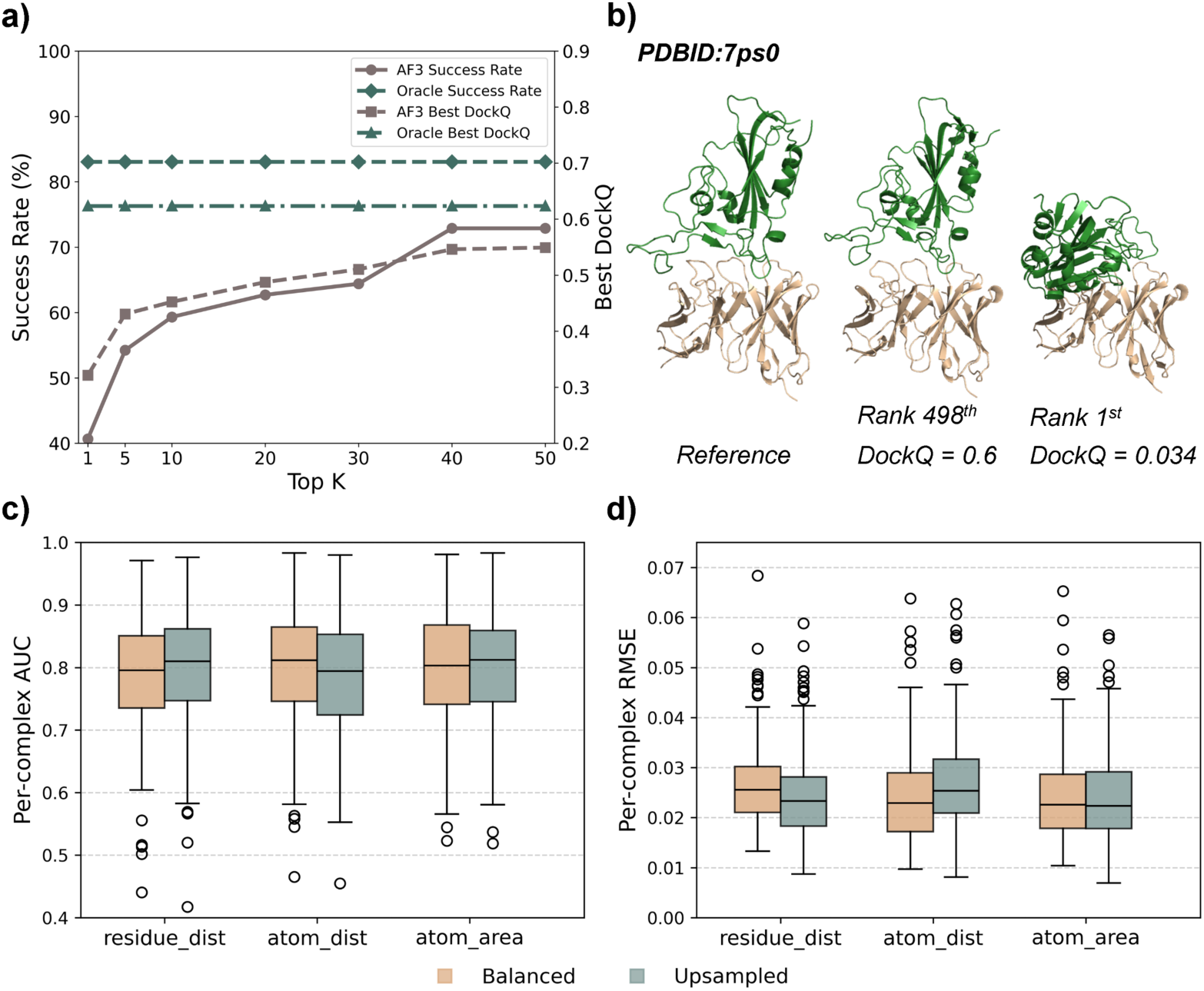
Performance Evaluation of AF3 and graph-based scoring algorithms. **a)** Evaluation of AF3’s scoring performance on 59 antibody-antigen complexes in our test set. The Top K success rate, where K represents the number of models considered, and Top K DockQ values at each K are shown. The “oracle” curve represents the ideal performance obtained when the best possible model is selected for each complex. Top K success rate is defined by counting the number of docking cases in which at least one near-native model is found among the Top K ranking models, divided by the total number of cases. Top K DockQ is computed by selecting the highest DockQ value among Top K ranked models for each complex and then averaging across complexes. **b)** One example where AF3 produces high quality models but ranks it 498th, while the Top 1 ranked model is incorrect. **c-d)** Comparison of model performance across graph representations and sampling strategies. Per-complex AUC (c) and per-complex MSE (d) for models trained using residue-level or atom-level graphs under “*balanced”* or “*upsampled”* sampling methodology (see Methods). “*residue_dist*” corresponds to residue-level graphs with distance-based edges; “*atom_dist*” and “*atom_area*” denote atom-level graphs using Euclidean distance or Voronoi contact area, respectively. Boxplots show the distribution of values across complexes averaged over five-fold cross-validations.

### 2.2 Design and optimizing DeepRank-Ab in terms of graph level, features, and sampling strategies

Having identified the scoring limitations of AF3, we developed DeepRank-Ab guided by two key principles: (i) constructing reliable and diverse training data, and (ii) identifying an effective structural representation for learning. To this end, we curated 1,442 antibody-antigen complexes from SAbDab and generated a new docking benchmark with HADDOCK3 ^25^. Details are provided in the Methods section.

Docking benchmarks are often redundant, raising a question of how to best leverage such large datasets. We therefore focused on two rarely discussed but critical design choices: (i) the level of structural information encoded in data representation, and (ii) the strategy to sample models from the benchmark dataset for training and testing.

### 2.3 Graph design

Graph-based learning has proven effective across many structural biology applications ^26,27^. For protein-protein interactions, previously we used residue-level graph representations for the scoring task. However, because antibody-antigen interactions rely on CDR loops that are highly flexible and structurally complex ^28^, we therefore examined whether representing structures at the atomic level could improve performance.

We further expanded the feature set to enrich the graph representation, adding CDR region labels for nodes feature; edge geometry and energetics, as well as atom-atom contact areas derived from Voronoi tessellation ^29^ as edge features. Full descriptions of these features are provided in the Methods section.

### 2.4 Sampling strategies for training and evaluation sets

A major challenge in training scoring functions is the distribution mismatch between training data and real docking outputs, where near-native decoys are rare and conformational changes are often observed. How to best sample decoys from docking benchmarks for efficient training remains an underexplored area in the field. Here, we considered two complementary motivations: first, balanced sampling, which stabilizes training process by ensuring a more even representation of decoy qualities. The second is targeted upsampling of low-quality decoys (DockQ ≤ 0.23), based on the observation that near-native models tend to resemble one another, whereas low-quality decoys, although more abundant, can fail in various ways. To systematically evaluate these considerations, we compared two sampling strategies: (i) a balanced sampling approach and (ii) targeted upsampling of low-quality (DockQ ≤ 0.23) decoys. The overview of our DeepRank-Ab workflow is shown in Figure 2. Refer to the Methods section for details of the sampling method.

**Figure 2.**
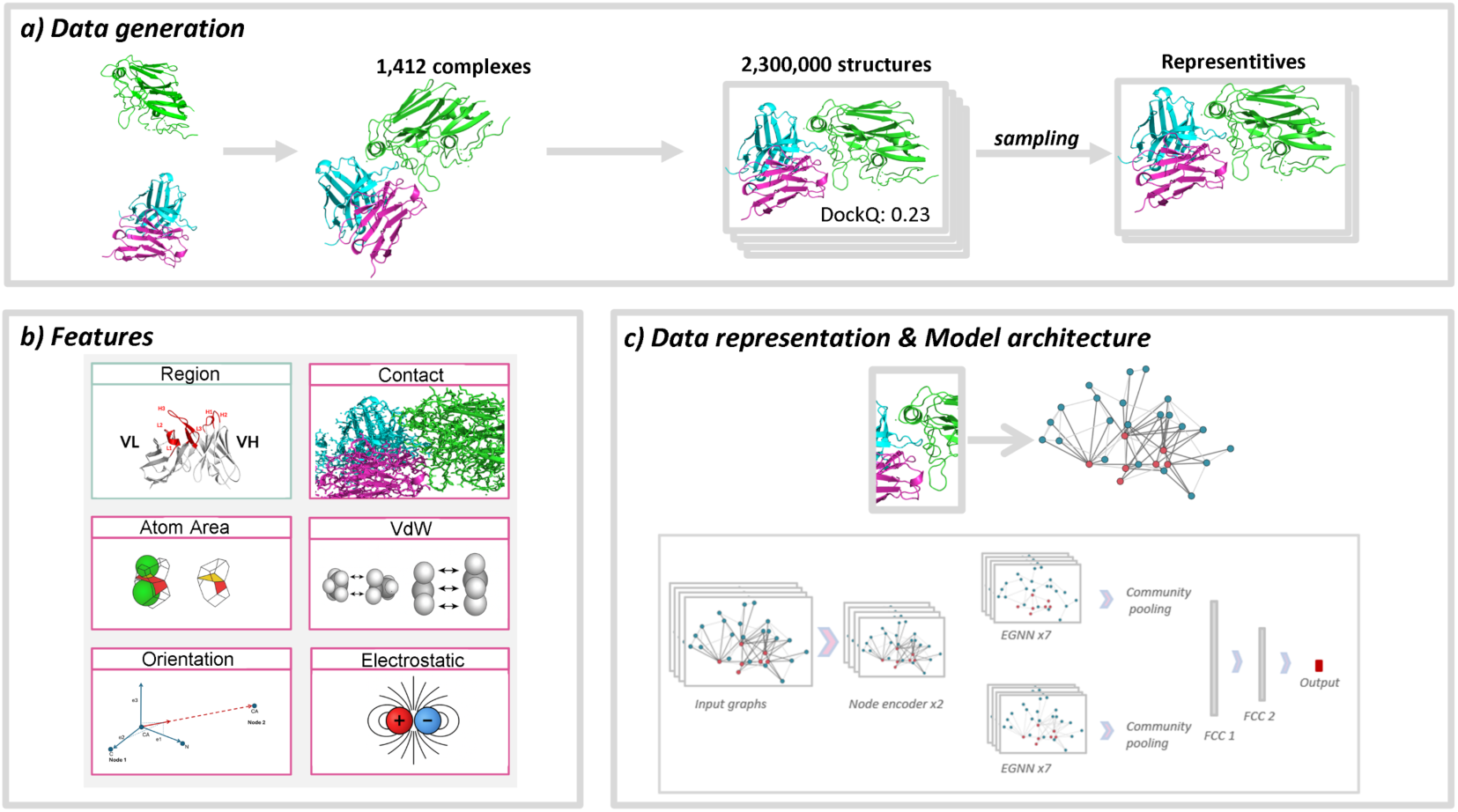
The various stages of the DeepRank-Ab workflow. **a)** Data generation: experimental structures were re-docked in different scenarios, resulting in 2.3 million models, from which representative datasets were sampled. **b)** Feature definition: new antibody-specific and physics-based descriptors were added to the existing features in DeepRank-GNN-esm. **c)** Interfaces were represented as graphs and used as inputs to train equivariant graph neural networks.

### 2.5 Model training and evaluation

Using combinations of the two sampling strategies and the two graph levels discussed as well as feature combinations that we discussed above, we trained six DeepRank-Ab variants and evaluated them using five-fold cross-validation. We report per-complex statistics because, in antibody-antigen scoring, overall AUC values can appear high even when performance on individual complexes is poor; per-complex evaluation is therefore more informative ^12^. Figure 1c-d summarizes the average per-complex MSE and AUC for each model across the five folds. Complementing the per-complex analysis, Table 1 reports the overall regression and binary classification performance of each DeepRank-Ab variant.

**Table 1.** Overall performance of DeepRank-Ab variants across five-fold cross-validation on benchmark evaluation set. This table reports the global regression and binary classification metrics (mean ± standard deviation) for all six DeepRank-Ab variants. Models differ in graph representation (residue- or atom-level), interaction encoding (Euclidean distance vs. Voronoi contact area), and sampling strategy (balanced vs. upsampled). Metrics include RMSE, R², AUC, Accuracy, Precision, and Recall, computed across all complexes in the dataset during evaluation.

|  | DeepRank-Ab variants |  |  |  |  |  |
| --- | --- | --- | --- | --- | --- | --- |
|  | *residue_dist* | *atom_dist* | *atom_area* | *residue_dist* | *atom_dist* | *atom_area* |
| Sampling | balanced | balanced | balanced | upsampled | upsampled | upsampled |
| RMSE (mean ± *std*) | 0.165 ± 0.002 | 0.165 ± 0.004 | 0.164 ± 0.003 | 0.157 ± 0.003 | 0.163 ± 0.005 | **0.155 ± 0.005** |
| $R^2$ (mean ± *std*) | 0.578 ± 0.011 | 0.575 ± 0.018 | **0.583 ± 0.012** | 0.508 ± 0.018 | 0.468 ± 0.035 | 0.518 ± 0.026 |
| AUC (mean ± *std*) | 0.784 ± 0.012 | 0.786 ± 0.014 | 0.787 ± 0.010 | 0.803 ± 0.011 | 0.794 ± 0.017 | **0.808 ± 0.016** |
| Accuracy (mean ± *std*) | 0.705 ± 0.010 | 0.706 ± 0.012 | 0.708 ± 0.009 | 0.777 ± 0.005 | 0.768 ± 0.011 | **0.784 ± 0.007** |
| Precision (mean ± *std*) | 0.750 ± 0.010 | **0.762 ± 0.013** | 0.757 ± 0.009 | 0.641 ± 0.013 | 0.622 ± 0.023 | 0.671 ± 0.014 |
| Recall (mean ± *std*) | **0.710 ± 0.013** | 0.690 ± 0.019 | 0.704 ± 0.029 | 0.542 ± 0.026 | 0.534 ± 0.021 | 0.513 ± 0.028 |

Four variants used atom-level graphs and differed in how interatomic interactions are encoded, either by Euclidean distance (‘*atom_dist’*) or by Voronoi contact area (‘*atom_area’*), and in their sampling strategy (‘*balanced’* vs. ‘*upsampled’*). Motivated by the strong performance of Voronoi-based scoring approaches ^17,30^, we examined explicitly how the area-based representations compare to distance-based alternatives.

As shown in Figure 1c-d, the Voronoi-based ‘*atom_area’* representation performs slightly better than ‘*atom_distance’ representation*, achieving lower or equal MSE under both sampling strategies (0.025 vs. 0.027 for ‘*upsampled*’ sampling; 0.025 vs 0.025 for ‘*balanced*’ sampling) and higher AUC or equal (0.801 vs. 0.785 for ‘*upsampled*’ sampling; 0.799 vs 0.799 for for ‘*balanced*’ sampling). Overall, atom-level graphs outperform residue-level ones, indicating that additional structural detail improves accuracy. The ‘*upsampled*’ variants show slightly better results, suggesting that enriching high-quality and diverse low-quality decoys benefits training.

### 2.6 Ablation study and feature importance analysis

To quantify the importance of individual features, we performed ablation studies on the best model (‘*upsampled atom_area’*) using five-fold cross-validation. Additional ablation experiments were also conducted on the full dataset Figure 3a demonstrates the impact of removing each feature in turn, reported as ΔRMSE relative to the baseline model (RMSE = 0.159). Removing region labels, atom type, covalent interactions, orientation, or Voronoi contact area caused the largest increases in RMSE and well as a larger variance across training folds, indicating that these features are essential for both accuracy and training stability. In contrast, removing embeddings, residue type, or charge caused only minor performance changes, suggesting partial redundancy. Ablations on the full dataset yielded similar trends (SI Figure 2). Interestingly, fine-tuned embeddings ^31^ improved RMSE, whereas raw ESM-2 embeddings did not, indicating that antibody-specific fine-tuning is necessary.

**Figure 3.**
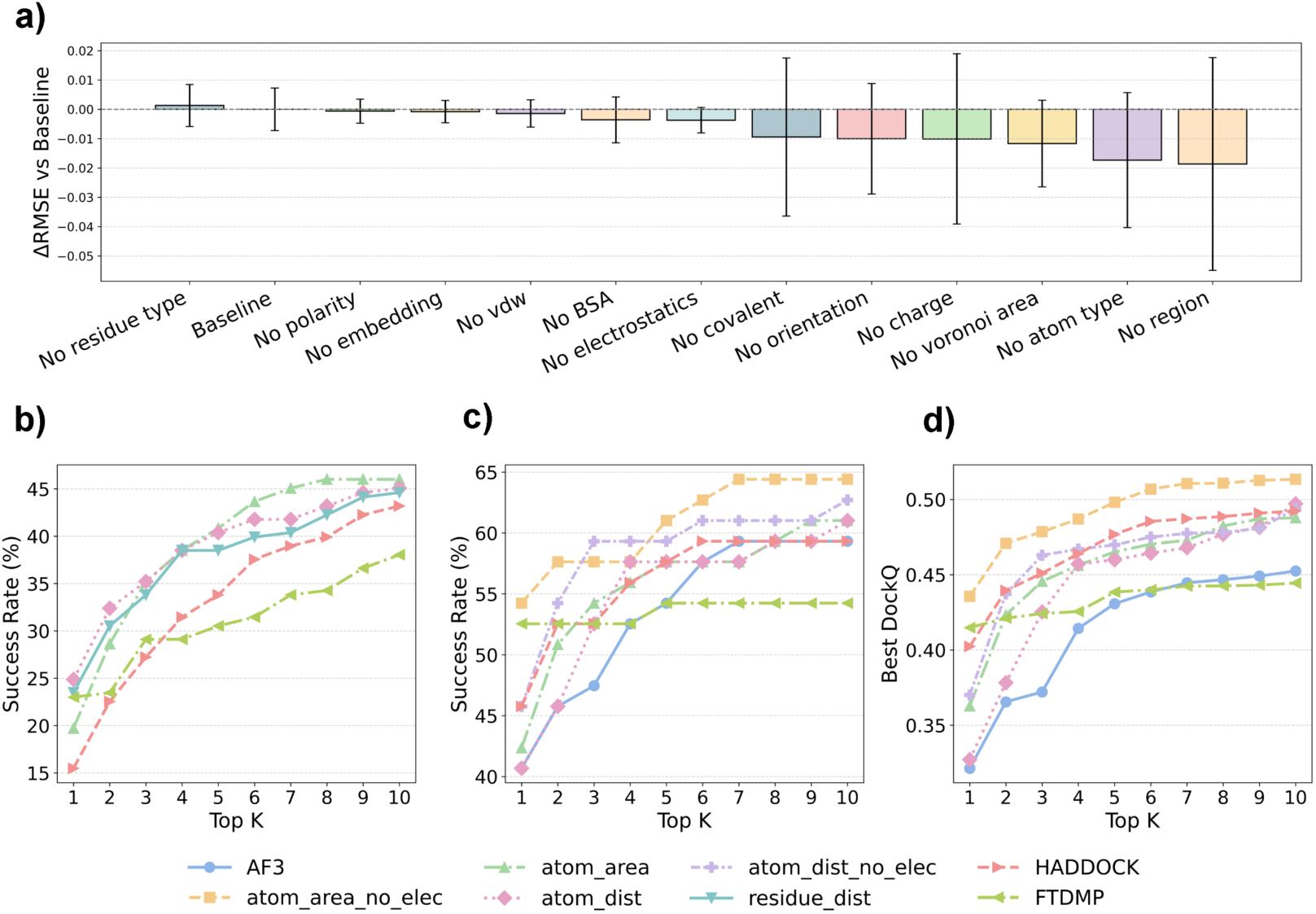
**a)** Ablation study (5-fold cross-validation). We report the mean RMSE ± SD obtained when each feature group is removed, relative to the full baseline model. **b)** Top k success rate of 5 methods on 215 complexes in benchmark test set, including 3 DeepRank-Ab variants (“*atom_area”, “atmo_distanc”,” residue_distance”*), HADDOCK EMscoring and FTDMP. **c-d)** Performance comparison of eight methods on 59 complexes from the AF3 test set, including AF3, three DeepRank-Ab variants with electrostatic features (“*atom_area*”, “*atom_distance*”, “*residue_distance*”), two variants without electrostatics (“*atom_area_no_elec*”, “*atom_distance_no_elec*”), HADDOCK EMScoring, and FTDMP. Panel c shows the Top K success rate; panel d shows the best Top K DockQ. Refer to SI Methods 5 for a detailed description of the feature set used by each model.

### 2.7 Performance on the test sets: benchmark test sets and AF3 test sets

**2.7.1 Performance on benchmark test sets**

To rigorously assess model performance under realistic docking conditions, we restricted our test set to the most challenging scenario: unbound unbound docking. Including models from the other three docking conditions would greatly inflate the Top 1 success rate because, in our experience, high-quality structures, particularly those produced by the HADDOCK refinement protocol, are very easy to rank.

We retrained the final models incorporating insights from the ablation study and compared them against the HADDOCK score obtained using the HADDOCK3 emscoring module and FTDMP ^17^. In Figure 3b, all three DeepRank-Ab variants outperform these baselines. The ‘*atom_dist’* model yields the highest Top 1 success rate, whereas ‘*atom_area’* performs best at Top 10. Classification and regression metrics of these DeepRank-Ab variants are also reported at Table 2.

**Table 2.** Overall performance of three DeepRank-Ab variants on benchmark test sets. This table reports the regression, binary classification metrics as well as Top K success rate (%). Variants differ in graph representation (residue- or atom-level), interaction encoding (Euclidean distance vs. Voronoi contact area).

| Graph representation | Regression Metrics |  |  | Classification Metrics |  |  | Success Rate (%) |  |  |
| --- | --- | --- | --- | --- | --- | --- | --- | --- | --- |
|  | RMSE | R<sup>2</sup> | AUC | Accuracy | Precision | Recall | T1 | T5 | T10 |
| residue_dist | 0.156 | 0.525 | 0.808 | 0.786 | 0.641 | 0.544 | 23.47 | 38.5 | 44.6 |
| atom_dist | **0.155** | **0.527** | **0.812** | **0.795** | **0.664** | 0.549 | **24.88** | 40.38 | 45.07 |
| atom_area | 0.16 | 0.498 | 0.79 | 0.769 | 0.594 | **0.564** | 19.72 | **40.85** | **46.01** |

We further investigated whether DeepRank-Ab’s ranking performance was associated with structural characteristics of the complexes, such as interface size or the lengths of key CDR loops (CDR-H1 and CDR-H3). However, none of these properties showed a meaningful correlation with the model’s ability to identify near-native solutions. In contrast, antigen length showed a strong effect: DeepRank-Ab ranked peptide antigens far more accurately than protein antigens in the test set (Top 1 success rate 52.38% vs. 11.56%, Top 5 success rate 88.09% vs 28.9%). Here, peptide antigens are defined as antigens with fewer than 50 residues, following the convention used in SAbDab ^32^. Using this definition, the 215 complex test set contains 42 peptide-antigen complexes.

Several factors likely contribute to the superior performance on peptides. First, antibody-peptide interfaces are known to have greater hydrophobic burial and higher shape complementarity ^33^, which may make them easier for the model to predict. Second, our graph construction uses a relatively small distance cutoff, focusing on local atomic interactions at the interface. This design choice is particularly well suited for peptide antigens, which are short and whose binding is dominated by local contacts rather than long-range interactions. In contrast, protein antigens often involve more extended interfaces which are more challenging for graph-based learning approaches to capture. Collectively, the combination of favourable interface properties and a graph representation tailored to local interactions likely explains the strong performance of our method on antibody-peptide complexes.

Although protein language-model embeddings showed only modest improvements in RMSE in our ablation study, they provided clear benefits for our primary objective—accurate ranking of top models rather than precise prediction of every DockQ value. However, we did not observe further gains when using fine-tuned embeddings.

#### 2.7.1 Performance on AF3 test sets

We next evaluated whether scoring functions trained on HADDOCK decoys generalize to AF3-generated structures. All tested methods outperform AF3 ranking score on the AF3 test set (Figure 3c). The best model (‘*atom_area*’) achieves a Top 1 success rate of 49.15% (vs. AF3 40.67% and HADDOCK 45.7%).

During analysis, we also observed that unrelaxed AF3 structures often exhibited energetic features that differed from those in our training set, likely due to local steric clashes or suboptimal side-chain packing. To mitigate this distribution shift, we trained additional models in which the electrostatic energy term was removed (‘*atom_area_no_elec’* and ‘*atom_dist_no_elec’*). This adjustment further improved performance, increasing the Top 1 success rate from 49.15% to 54.24% (Figure 3. c), making DeepRank-Ab the best-performing method on this dataset.

Beyond success rates, DeepRank-Ab is also more effective at prioritizing high-quality structures. As shown in Figure 3d, the mean Top 1 DockQ increased by 35.5%, from 0.321 for the AF3 baseline to 0.435 with our best-performing method. In 99% of cases, when DeepRank-Ab correctly identified a Top 1 structure among the HADDOCK-generated models for a given complex, it also recovered the correct Top 1 model from AF3. This further illustrates the extent to which DeepRank-Ab generalizes across different model-generation protocols.

Finally, motivated by the success of the consensus scoring approach in previous rounds of CAPRI ^34^, we evaluated two consensus strategies: by averaging the model predictions and or by applying a jury-based voting method as in VoroIF-GNN ^30^, combining predictions from our best model with AF3. However, both strategies reduced performance (Table 3). Consensus scoring is most effective when the combined methods contribute complementary information. In our case, our top-performing model is already highly optimized, combining it with other methods dilutes rather than improves its ranking power.

**Table 3.** Comparison of regression metrics, classification metrics, and Top K success rates for DeepRank-Ab variants, consensus scoring strategies, and AF3’s native ranking score on 59 AF3-generated antibody-antigen complexes. Variants differ in graph representation (residue- or atom-level), interaction encoding (Euclidean distance vs. Voronoi contact area). ‘no_elec’ variant means the model variant trained without electrostatic features. The two consensus methods are: (1) consensus_jury, a jury-based voting scheme combining AF3 with our best model (“atom_area_no_elec”), and (2) consensus_average, which uses the averaged ranks from AF3 and “atom_area_no_elec” as the final ranking

| Graph representation | Regression Metrics | Binary Classification Metrics |  |  |  | Success Rate (%) |  |  |
| --- | --- | --- | --- | --- | --- | --- | --- | --- |
|  | RMSE | AUC | Accuracy | Precision | Recall | T1 | T5 | T10 |
| residue_dist | **0.337** | 0.682 | **0.572** | **0.467** | 0.749 | 35.59 | 49.15 | 57.63 |
| atom_dist | 0.477 | 0.839 | 0.403 | 0.399 | **0.999** | 40.68 | 57.63 | 61.02 |
| atom_area | 0.518 | 0.851 | 0.4 | 0.398 | **0.999** | 42.37 | 57.63 | 61.02 |
| atom_dist_no_elec | 0.431 | 0.786 | 0.537 | 0.459 | 0.951 | 45.76 | 59.32 | 62.71 |
| atom_area_no_elec | 0.399 | **0.853** | 0.542 | 0.464 | 0.978 | **54.24** | **61.02** | **64.41** |
| consensus_jury |  |  |  |  |  | 37.28 | 45.76 | 54.24 |
| consensus_average |  |  | N/A |  |  | 50.84 | **61.02** | 61.02 |
| AF3 |  |  |  |  |  | 40.68 | 54.24 | 59.32 |

To further assess DeepRank-Ab, we evaluated it on an external dataset of five MassiveFold ^35^ targets of antibody-antigen complexes taken from recent rounds of CAPRI ((Lensink et al.). Four of these targets involve peptide antigens. DeepRank-Ab showed very strong performance on this dataset, achieving a 100% success rate at Top 5, while AF3 reached only 60% (SI Figure 3). These results are consistent with our earlier observations that DeepRank-Ab is particularly effective for peptide-antigen complexes.

## 3. Methods

### 3.1 Dataset construction and generation of the docking benchmark

We constructed a non-redundant dataset of antibody-antigen complexes from SAbDab^32^ by first retrieving all eligible structures and then applying several filtering steps. Concatenated CDR similarity filtering and structural quality thresholds (resolution ≤ 3.5 Å; R-factor ≤ 0.25) were used to remove redundant or low-quality entries, along with additional checks to exclude incomplete antibodies and non-protein antigens. For antigens composed of multiple chains, only those in contact with the antibody were retained. To improve the modelling efficiency, antigens longer than 500 residues were segmented using Merizo ^36^, and only high-confidence domains (score > 0.7) containing interface residues were kept. Entries for which ABodyBuilder2 ^37^ failed to generate antibody models or for which AF2 yielded poor antigen predictions (RMSD > 8 Å) were removed. After these steps, the final curated dataset comprised 1,442 antibody-antigen complexes, which served as input to HADDOCK3 ^25^.

When generating the docking benchmark, we had several design objectives. First, we aimed to obtain a diverse set of low-quality (DockQ ≤ 0.23) models while also ensuring that each complex contained at least one high-quality model. Conformational diversity was introduced through a combination of docking and refinement procedures. To achieve this, we applied four HADDOCK3 protocols, including two docking scenarios (bound-bound docking and unbound-unbound docking) and two refinement scenarios (bound refinement and unbound refinement). Details are provided in the SI Methods 1 section.

To construct representative decoy sets for training, we applied two sampling strategies. In refinement-only scenarios, all generated models were retained. In docking scenarios, decoys from rigid-body docking and energy minimization were grouped by DockQ score into bins of width 0.1, clustered, and a fixed number of models was selected from each bin and scenario (‘*balanced*’ sampling). In the second strategy (‘u*psampled*’), low-quality decoys (i.e., those in bin 0 and bin 1) were upsampled by a factor of 20 (See SI Methods 2, sampling statistics in SI Figure 1), leveraging HADDOCK’s wide conformational search to introduce additional structural diversity and improve training robustness.

### 3.2 Strict structural splitting to avoid data leakage

To prevent data leakage during training, all complexes were first clustered with Foldseek-Multimer ^38^ with thresholds of 0.65 for multimer TM, 0.5 for chain-level TM, and 0.65 for interface lDDT, defining interfaces with an 8.5 Å cutoff. Results showed that 89% of complexes end up in their own cluster, indicating adequate diversity in the dataset. Approximately 10% were grouped into clusters of size 2-9, and only 0.3% belonged to larger clusters (See SI Table 1 for clustering statistics). The dataset was then partitioned at the cluster level: clusters were randomly ordered and assigned to the training/validation split until ∼85% of all complexes were included, with the remaining ∼15% allocated to the test set. All decoys generated from a given complex were assigned to the same split as their parent complex.

### 3.3 Graph representation, new features, and model architecture

We formulated the scoring task as a regression problem in which a docking model is given as input and the network predicts an interface-quality score (pDockQ). Antibody-antigen interfaces were encoded as graphs at two levels of details. First, we constructed residue-level graphs, with nodes and edges defined following our previous DeepRank-GNN-esm framework ^18^. Second, we built atom-level graphs, in which heavy atoms serve as nodes and edges are defined either by interatomic distances or by contact areas computed using Voronoi tessellation ^29^. To enrich these representations, we introduced several new node- and edge-level features: nodes were annotated with antibody-specific region labels derived from IMGT numbering ^39^, and edges were assigned geometric descriptors capturing local residue orientation, along with energetic and covalent interaction terms adapted from DeepRank2 ^40^.

For the model architecture, we adopted an E(n)-equivariant GNN ^41^, chosen for its ability to incorporate geometric information while preserving rotational and translational symmetries. Following the design principles of DeepRank-GNN ^19^, we implemented a two-branch architecture that processes interface edges and internal edges separately, with each branch consisting of seven EGNN layers. Edge messages integrate node features, RBF-encoded distances, and the full set of edge attributes, while coordinate updates are computed via learned transformations. Node updates employ residual connections, LayerNorm, and dropout. After message passing, we apply hierarchical pooling followed by a Global Attention readout. The resulting embeddings from both branches are concatenated and passed through a lightweight prediction head with a residual connection to generate the final pDockQ score. Full description of graph design, feature implementation, and training details see SI Methods sections 3 and 4.

### 3.4 Curation of external test sets

We evaluated our method on three test sets: our own Benchmark test set, the AF3 test set, and five MassiveFold ^35^ targets from previous round of CAPRI. For the Benchmark test set, we included only unbound-unbound docking models with upsampled acceptable, lower-quality models. For the AF3 test set, we generated 500 structures per complex using a local installation of AF3 ^4^ with 100 random seeds. 5 CAPRI targets were taken as external test sets (T231, T234, T255, T268, T270), with an average of 8,046 models generated with MassiveFold ^35^.

## 4. Conclusions

In this work, we addressed the scoring challenge for antibody-antigen complexes and showed that existing methods, including AF3, have substantial limitations in their ability to rank near-native structures. To support the development of DeepRank-Ab, we curated a high-quality and structurally diverse docking benchmark specifically tailored to antibody-antigen interactions. This dataset was generated using multiple HADDOCK3 workflows, including both docking and refinement protocols, combined with a sampling strategy designed to maximize structural diversity.

Using this dataset, we conducted a systematic evaluation of graph representations and feature sets and developed DeepRank-Ab, including a suite of scoring functions optimized for structures produced by both docking pipelines and AF3. Across multiple independent test sets, DeepRank-Ab consistently demonstrated superior performance, achieving the highest Top k success rates and Top k DockQ values compared with AF3, HADDOCK scoring, and FTDMP.

We anticipate that supplementing AF3 with DeepRank-Ab should compensate, at least in part, for the absence of explicitly incorrect structures in AF3’s training regime. Although AF3 is indirectly exposed to suboptimal conformations during its recycling procedure and possibly through entries in AlphaFoldDB ^4^, systemic datasets such as docking decoys may still introduce additional types of structural variation that are not represented in its training data.

## Author contributions

X.X, I.C and A.M.J.J.B designed the research. X.X and I.C conducted the research and analysed the results. I.C generated the HADDOCK3 docking benchmark. X.X and I.C contributed to the graph design, training and evaluation of model architecture. V.R contributed to graph featurization. All authors wrote and reviewed the manuscript.

## Acknowledgements

We acknowledge Marc Lensink and Guillaume Brysbaert, University of Lille, France, for providing the 5 MassiveFold targets.

## Data and code availability

All data involved in this work is available at https://zenodo.org/10.5281/zenodo.17911452. DeepRank-Ab is available online at https://github.com/haddocking/DeepRank-Ab.

