## Supplementary material for "DeepRank-Ab: a scoring function for antibody-antigen complexes based on geometric deep learning"

**Table S1. Antibody-antigen complex cluster size distribution shown as both percentage and absolute number of structures.**

| Cluster Size | Percentage (%) | Number of complexes |
|---|---|---|
| 1 | 89.19 | 1031 |
| 2 | 7.27 | 84 |
| 3 | 1.82 | 21 |
| 4 | 0.69 | 8 |
| 5 | 0.35 | 4 |
| 6 | 0.17 | 2 |
| 7 | 0.09 | 1 |
| 9 | 0.09 | 1 |
| 16 | 0.09 | 1 |
| 28 | 0.09 | 1 |
| 56 | 0.09 | 1 |

**Table S2. Summary of all model variants evaluated in this study, including their sampling strategies, graph representations, and associated feature sets.** Models in Figure 1c-d compare residue-level and atom-level graphs trained with either balanced or up-sampled sampling. Residue-level models ("*residue_dist*") use residue-based node features and distance-based edges, whereas atom-level models ("*atom_dist*" and "*atom_area*") incorporate atom-specific features with either Euclidean distance or Voronoi contact-area edges. Models used in Figure 3 explore extended and reduced feature sets as well as the impact of removing electrostatic edge features ("*no_elec*"). All Figure 3 models employ atom-level graphs and up-sampled training.

| Related Figure | Model Name | Sampling Method | Graph level | Node features | Edge features |
|---|---|---|---|---|---|
| Figure 1. cd | residue_dist | Balanced sampling | residue | [polarity,bsa,charge,region,embedding,type] | [dist,covalent,elec,vdw,orientation] |
| Figure 1. cd | atom_dist | Balanced sampling | atom | [atom_type,polarity,bsa,charge,region,embedding,res_type] | [dist,covalent,elec,vdw,orientation] |
| Figure 1. cd | atom_area | Balanced sampling | atom | [atom_type,polarity,bsa,charge,region,embedding,res_type] | [voro_area,covalent,elec,vdw,orientation] |
| Figure 1. cd | residue_dist | Up-sampled | residue | [polarity,bsa,charge,region,embedding,type] | [dist,covalent,elec,vdw,orientation] |
| Figure 1. cd | atom_dist | Up-sampled | atom | [atom_type,polarity,bsa,charge,region,embedding,res_type] | [dist,covalent,elec,vdw,orientation] |
| Figure 1. cd | atom_area | Up-sampled | atom | [atom_type,polarity,bsa,charge,region,embedding,res_type] | [voro_area,covalent,elec,vdw,orientation] |
| Figure 3. a | atom_area | Up-sampled | atom | [atom_type,polarity,bsa,charge,region,embedding,res_type,embedding_finetrue] | [voro_area,covalent,elec,vdw,orientation] |
| Figure 3. b | atom_area | Up-sampled | atom | [atom_type,polarity,bsa,region,embedding] | [voro_area,covalent,elec,vdw,orientation] |
| Figure 3. b | atom_dist | Up-sampled | atom | [atom_type,polarity,bsa,region,embedding] | [dist,covalent,elec,vdw,orientation] |
| Figure 3. cd | atom_area_no_elec | Up-sampled | atom | [atom_type,polarity,bsa,region,embedding] | [voro_area,covalent,vdw,orientation] |
| Figure 3. cd | atom_dist_no_elec | Up-sampled | atom | [atom_type,polarity,bsa,region,embedding] | [dist,covalent,vdw,orientation] |

**Figure S1. Statistics of two sampling methods. a)** Global distribution of DockQ bins in our benchmark across all 1,442 complexes. **b)** Percentage of 'Acceptable' and 'Incorrect' models (Acceptable defined as DockQ ⩾ 0.23).

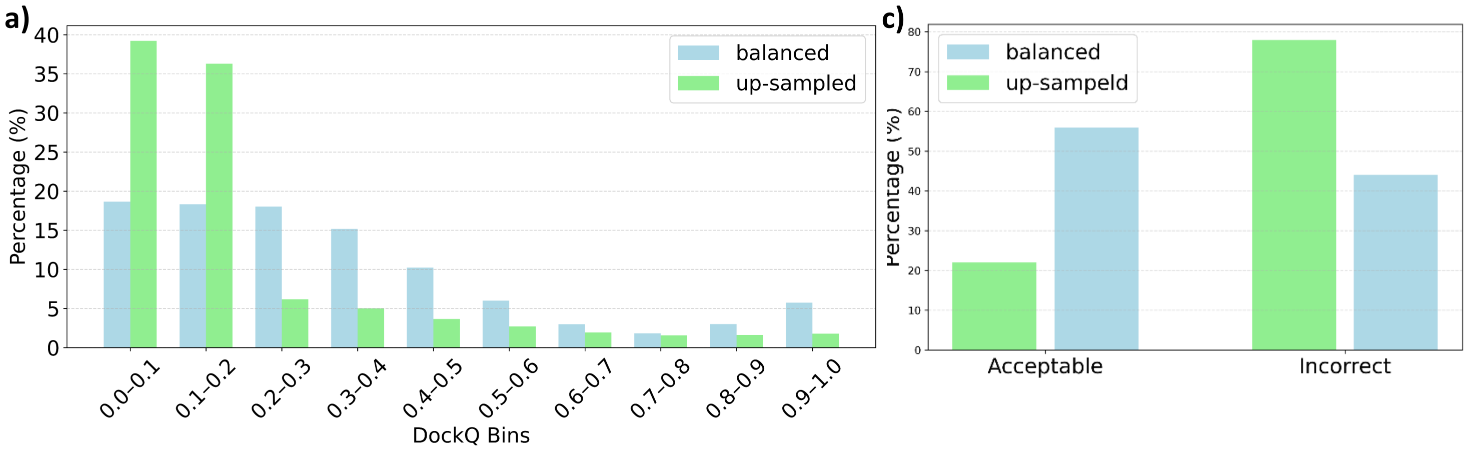

**Figure S2. Ablation study on the full training set**

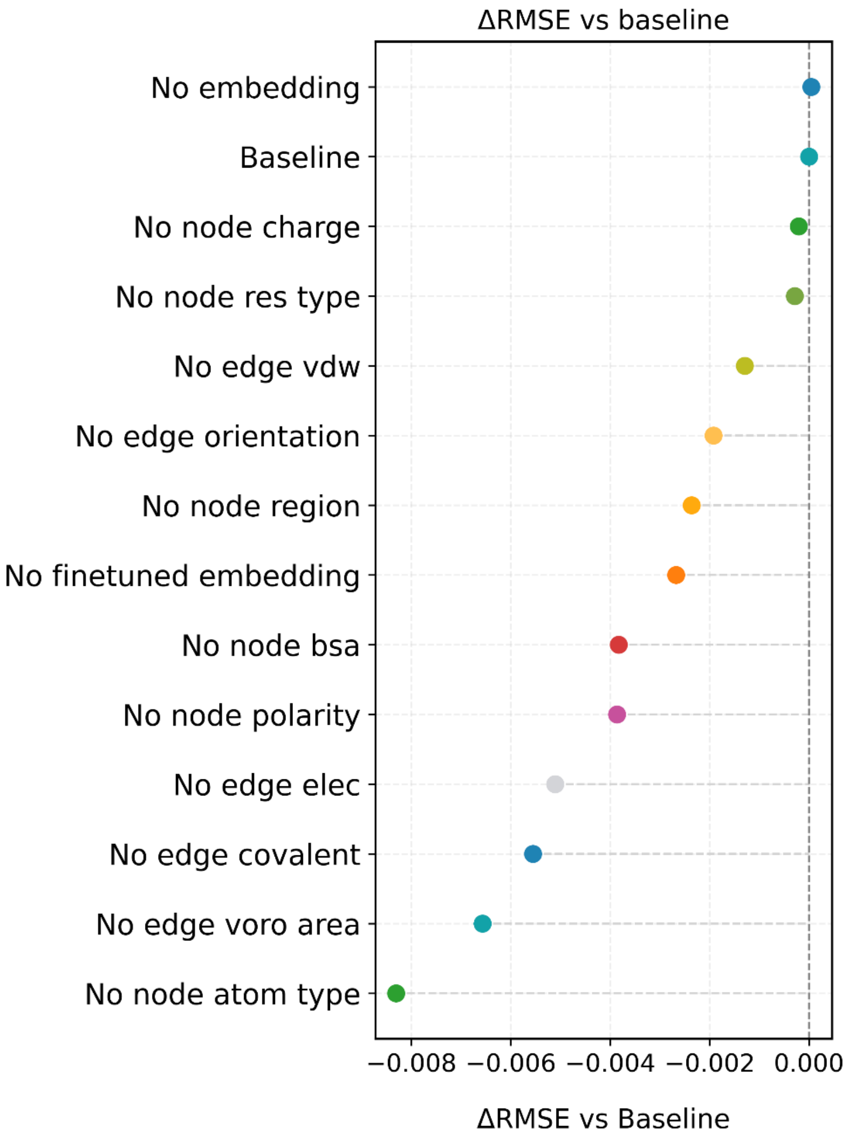

**Figure S3. Top K success rate and Top K DocKQ of DeepRank-Ab, AF3 and HADDOCK on 5 MassiveFold targets.**

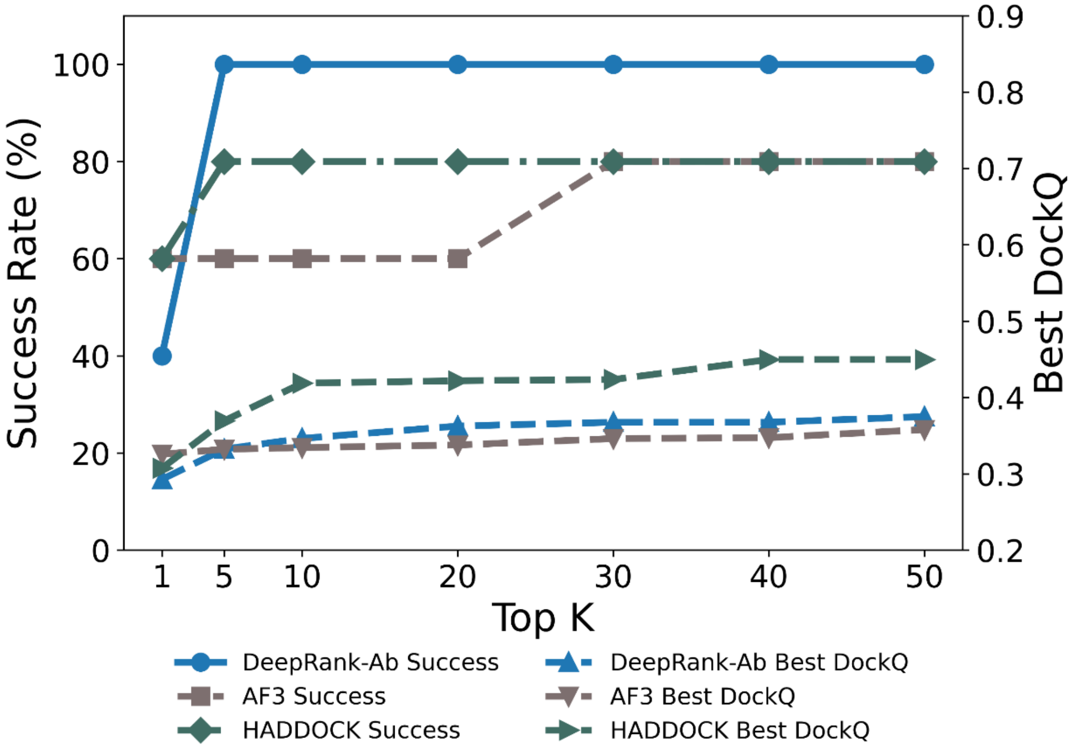

### Methods section S1. HADDOCK modelling workflows for the docking benchmark

To build the pool of docking decoys, we used the haddock-runner (https://www.bonvinlab.org/haddock-runner/) to execute HADDOCK3 (Giulini *et al.*, 2025) under four scenarios. These scenarios differed in whether bound or predicted structures were used and in the restraints applied during sampling. Across all scenarios, we generated a total of 2,278,533 decoys. Details as below:

- **scenario 0**: Bound Redock and Refine (redocking and then refining the experimental bound complexes);
- **scenario 1**: Bound Refine-only (refining the experimental complex without redocking);
- **scenario 2**: Predicted unbound Dock and Refine AI-generated models (ABodyBuilder2 (Abanades *et al.*, 2023) for antibodies, AlphaFold2 for antigens)
- **scenario 3**: Unbound Align and Refine (refining AI-generated unbound models after rigid alignment to the bound structures). In total, we generated 2,278,533 decoys.

In *Scenario 0* (Bound Redock + Refine), we split the 1,442 experimental complexes into antibody and antigen chains and applied the preprocessing steps above. The antibody is treated as a single chain by shifting the numbering of the light chain to avoid overlap (https://github.com/haddocking/pdb-tools) (Rodrigues *et al.*, 2018). The same applies to multi-chain antigens. For both the antibody and the antigen in each complex, we defined multi-bodies unambiguous restraints for both antibody and multi-chain antigen partners using *haddock3-restraints restrain_bodies*. These are used in HADDOCK to keep unconnected domains together during the flexible refinement (e.g. the heavy and light chains of the antibody). Ambiguous interactions restraints between the antibody and antigen were generated using a 5 Å distance cutoff using the *haddock-restraints ti* command (https://github.com/haddocking/haddock-restraints). These restraints were combined. The orientation of the input structures is randomized for the rigid body docking stage. *Scenario 1* (Bound Refine-only) reused the input structures from Scenario 0 to maintain consistency across bound cases but keeps those in their original positions (i.e. the orientation in the complex). For the refinement, only multi-bodies unambiguous restraints were used.

In *Scenario 2* (Predicted Unbound Dock + Refine), we used AI-generated models instead of experimental structures. Antibodies were trimmed to their variable domains, cleaned as above, and their sequences were extracted and submitted to ABodyBuilder2. The resulting light- and heavy-chain models were merged and renumbered as a single antibody chain. Antigens were similarly cleaned, their sequences extracted, and predicted using AlphaFold2 (alphafold2_ptm for monomers and alphafold2_multimer_v3 for multimers) before merging and renumbering as chain B. From these models, we constructed multi-bodies unambiguous restraints as in Scenario 0. Ambiguous restraints were built by designating the antibody CDRs as active residues (relative solvent accessibility threshold of 0.15) and antigen residues within 50 Å of the antibody as passive residues (relative accessibility threshold 0.3), *using haddock3-restraints calc_accessibility* followed by *active_passive_to_ambig*.

*Scenario 3* (Predicted Unbound Align + Refine) also relied on the predicted antibody and antigen models generated in Scenario 2, but aligned onto the experimental complexes using TM-align. After alignment, models were cleaned and renumbered, and only multi-bodies unambiguous restraints were applied for refinement.

We executed all docking workflows through the *haddock-runner*. Scenarios 0 and 2 followed a full restrained docking pipeline, consisting of the modules *topoaa*, *rigidbody*, *clustfcc*, *seletopclusts*, *caprieval*, *flexref*, *emref*, and *emscoring*, with a tolerance for failure set to 20%. Rigid-body sampling generated 10,000 structures per complex, using both unambiguous and ambiguous restraints. After rigid docking, we performed clustering with *clustfcc*, selected up to 100 clusters, and retained the five top-scoring models from each cluster (maximum 500 structures). These were refined through flexible refinement and energy minimization, and evaluated with CAPRI metrics and per-interface energy scoring. Scenarios 1 and 3 used a streamlined refinement-only workflow consisting of *topoaa*, *flexref*, *emref*, *caprieval*, and *emscoring*, also with a failure tolerance of 20. Note that this differs from the usual refinement scenario of HADDOCK as a flexible refinement step (*flexref*) consisting of a simulated annealing in torsion angle space is added to the protocol to allow the models to deviate a bit more from the original structures. The final *emscoring* module was added to extract the per-interfae scores. Only multi-bodies unambiguous restraints were imposed, and one refined structure was produced per input model before evaluation with CAPRI metrics. Example HADDOCK3 config files for each scenario are available at [link].

#### Methods 2: Decoy sampling procedure

We developed a standardized procedure to construct representative decoy sets across docking scenarios. For Scenarios 1 and 3, no additional filtering was required because each complex yielded only a single refined structure. In contrast, Scenarios 0 and 2 produced large and diverse decoy pools, motivating the use of two alternative sampling strategies. In both strategies, we first separated the docking outputs into two categories—rigid-body docking models (*rigidbody*) and refined models after energy minimization (*emref*). The selection procedure was then applied independently to each category and scenario, generating four parallel processing streams: Scenario 0-*rigidbody*, Scenario 0-*emref*, Scenario 2-*rigidbody*, and Scenario 2-*emref*.

In the first strategy ("balanced"), decoys from the docking scenarios were grouped by DockQ score into bins of width 0.1 (0.0-0.1, …, 0.9-1.0). Within each DockQ bin, structural redundancy was reduced using *Foldseek easy-multimercluster* with multimer TM-score ≥ 0.65, chain-level TM-score ≥ 0.5, and interface lDDT ≥ 0.65 (default parameters). From each bin and for each docking category, we aimed to select up to 15 representative decoys—5 from Scenario 0 and 10 from Scenario 2—allowing at most one model per cluster. In this strategy, if fewer than 15 models were available after inter-scenario balancing, we added additional models from already-used clusters within the same scenario, and finally, if needed, from the opposite scenario until the bin was filled or no more models were available. Because the procedure was run independently for *rigidbody* and *emref*, each DockQ bin could contribute up to 30 decoys in total (15 from rigidbody and 15 from emref), subject to availability. The second strategy ("upsampled") followed the same binning and clustering procedure but increased the representation of low-quality decoys by upsampling bins 0 and 1 (DockQ < 0.2) by allowing up to 20 structures per cluster to be considered during selection while keeping the overall per-scenario quotas unchanged.

**Methods 3: Graph generation and feature design**

We represented each docking model as a directed graph using DeepRank-GNN-esm. Nodes correspond to either residues or atoms that participate in at least one inter-chain contact (within 5 Å between chains A and B). Edges are added in both directions: (i) interface edges connect residues or atoms across chains if their minimum inter-atomic distance (calculated over all pairs of atoms of the two residues) is below 5 Å, while (ii) internal edges connect residues or atoms within the same chain if they are both or belong to interface residues and their minimum inter-atomic distance is below 3 Å. For residue graphs, the distance associated with each edge is this minimum atom-atom distance, expressed in Å. For atom graphs, the strength of an edge is either defined by atom-atom distance or atom contact area (see definition below). We retained the feature set from DeepRank-GNN esm while adding new features.

In addition, we introduced three new groups of features: Region (a node feature specific to antibodies), geometric edge features, and contact edge features.

*1. Geometric features (edges):*

1) *Orientation.* This feature captures the direction from residue i to j in the local frame of the source residue, following the definition in OAGNN (Li *et al.*, 2025). For each residue *i*, we build an orthonormal basis anchored at *Cα*: the first axis points from *Cα* to *C*, the second comes from the *Cα-N* vector orthogonalized to the first, and the third is their cross product. We compute the unit direction between the two Cα atom:

$$\boldsymbol{d_{ij}} = |\boldsymbol{x_j^{CA}} - \boldsymbol{x_i^{CA}}|$$

and express it in the local basis $\boldsymbol{R_i} = [\boldsymbol{e_1}\ \boldsymbol{e_2}\ \boldsymbol{e_3}]$ as

$$\boldsymbol{o_{ij}} = \boldsymbol{R_i^T}\ \boldsymbol{d_{ij}}$$

2) *Atom area.* We rely on Voronota-LT (Olechnovič and Grudinin, 2025) to compute atom-atom interface areas using a Voronoi-based radical tessellation, in which bisector planes are defined by both atomic radii and the probe radius. This procedure assigns each atom a precise, geometry-derived contact area with its neighbours. For every interacting atom pair, the corresponding interface area is added as an edge feature in our graph representation.

*1. Energetic and covalent features (edges).*

We adopted the contact-feature pipeline from DeepRank2 (Crocioni *et al.*, 2024) that implements the OPLS force-field parameters taken from HADDOCK (Dominguez *et al.*, 2003). For every residue-residue edge (interface at 5 Å; internal at 3 Å), we retrieved all atoms with *pdb2sql* (Renaud and Geng, 2020) and built the full atom-atom distance matrix, setting self-distances to infinity. We then computed per-atom pair energies. We stored three scalar edge features:

1) *elec (Coulomb).* The electrostatic energy is calculated as

$$\boldsymbol{E}_{\text{elec}} = \frac{\boldsymbol{k}\,\boldsymbol{q}_i\boldsymbol{q}_j}{\boldsymbol{\varepsilon}_0\,\boldsymbol{r}_{ij}}$$

with $\boldsymbol{k}$ = 332.0636 and $\boldsymbol{\epsilon}_0$ = 1.0, using per-atom partial charges from DeepRank2. We normalize this feature using scikit-learn RobustScaler fitted once on all available edges in the dataset and then applied to both interface and internal edges.

2) *vdw (Lennard-Jones).* The van der Waals energy is calculated as

$$\boldsymbol{E}_{\text{vdw}} = 4\boldsymbol{\varepsilon}_{ij}\left[\left(\frac{\boldsymbol{\sigma}_{ij}}{\boldsymbol{r}_{ij}}\right)^{12} - \left(\frac{\boldsymbol{\sigma}_{ij}}{\boldsymbol{r}_{ij}}\right)^{6}\right]$$

Using the Lorentz-Berthelot mixing rules, the Lennard-Jones parameters for an atom pair are computed as $\boldsymbol{\sigma}_{ij} = \frac{\sigma_i+\sigma_j}{2}$ and $\boldsymbol{\varepsilon}_{ij} = \sqrt{\boldsymbol{\varepsilon}_i\boldsymbol{\varepsilon}_j}$. For atoms on the same chain and within 4.2 Å, we apply the specific 1-4 interaction parameters ($\boldsymbol{\sigma}_{14}$, $\boldsymbol{\varepsilon}_{14}$). For 1-3 interactions where the interatomic distance is below 3.6 Å, both the Coulomb and van der Waals terms are set to zero. No additional scaling is applied to this feature.

3) *covalent (binary*). A flag set to 1 when the minimum interatomic distance between the two residues is < 2.1 Å, and 0 otherwise.

*3. Embedding and fine-tuned embeddings (Node)*

We generated sequence embeddings using the same procedure as in DeepRank-GNN-esm (Xu and Bonvin, 2024). For fine-tuned embeddings, we applied our AbTune (Xu and Bonvin, 2025) pipeline, in which each antibody sequence was fine-tuned using the concatenated heavy- and light-chain sequence with 50% LoRA layers over 10 steps. After fine-tuning, we computed the averaged embedding across the model's embedding dimension so that each residue node is represented by a single embedding vector.

*4. Region (nodes)*

We included Region, a node feature specific to antibodies. For each complex, we annotated the antibody residues on chain A with IMGT regions using ANARCI (Dunbar and Deane, 2016). We aligned chain A to the heavy (H) and light (L) reference sequences available for that complex, recovered the variable domains, and numbered them with ANARCI (IMGT scheme, germline assignment on). We then mapped IMGT positions to six CDR labels (L1: 27-38, L2: 56-65, L3: 105-117; H1: 27-38, H2: 56-65, H3: 105 117). Positions within the variable domains but outside these ranges became framework (FR), and any antibody residue outside the variable domains became CONST. We stored the annotations per model and, when building the residue graph, we assigned each antibody node a one-hot Region vector over nine classes: FR, L1, L2, L3, H1, H2, H3, CONST, AG. We labelled all antigen residues on chain B as AG.

*5. Original features described in DeepRank-GNN-esm*

*1) Type*: Amino acid type represented using one-hot encoding.

*2) BSA*: Calculated buried surface area of the complex.

*3) Polarity*: Polarity was assigned to one of four classes—apolar, polar, negatively charged, or positively charged—and represented through one-hot encoding.

*4) Charge*: Charge was assigned based on Kyle and Doolittle hydrophobicity scale.

**Methods section S4. Training details of DeepRank-Ab**

We trained all models on an NVIDIA RTX 6000 Ada GPU. Optimization used Adam with learning rate $2\times10^{-3}$ and batch size 128 for 100 epochs. We used the Balanced MSE loss (BMC) with a learnable noise variance. After back propagation we applied gradient clipping with L2 norm 1.0. We maintained an EMA (Exponential Moving Average) of the weights ($\beta = 0.999$) with a warmup of about two epochs, and we used the EMA weights for validation and testing. Early stopping had a patience of 15 epochs. We used the same procedure for models and for all ablation variants reported in the Results section. For each model we performed 5-fold cross validation on the train+val split, using the cluster-aware partitioning described in Methods section to define the After completing cross-validation, we trained our final models on the full training dataset using the same protocol. These fully trained models were then used to evaluate performance on both the AF3 test set and the benchmark test set.
